# plantR: An R package and workflow for managing species records from biological collections

**DOI:** 10.1101/2021.04.06.437754

**Authors:** Renato A. F. de Lima, Andrea Sánchez-Tapia, Sara R. Mortara, Hans ter Steege, Marinez F. de Siqueira

## Abstract

1. Species records from biological collections are becoming increasingly available online. This unprecedented availability of records has largely supported recent studies in taxonomy, biogeography, macroecology, and biodiversity conservation. Biological collections vary in their documentation and notation standards, which have changed through time. For different reasons, neither collections nor data repositories perform the editing, formatting, and standardization of the data, leaving these tasks to the final users of the species records (e.g. taxonomists, ecologists and conservationists). These tasks are challenging, particularly when working with millions of records from hundreds of biological collections.
2. To help collection curators and final users perform those tasks, we introduce `plantR`, an open-source package that provides a comprehensive toolbox to manage species records from biological collections. The package is accompanied by the proposal of a reproducible workflow to manage this type of data in taxonomy, ecology, and biodiversity conservation. It is implemented in `R` and designed to handle relatively large data sets as fast as possible. Initially designed to handle plant species records, many of the `plantR` features also apply to other groups of organisms, given that the data structure is similar.
3. The `plantR` workflow includes tools to (1) download records from different data repositories, (2) standardize typical fields associated with species records, (3) validate the locality, geographical coordinates, taxonomic nomenclature, and species identifications, including the retrieval of duplicates across collections, and (4) summarize and export records, including the construction of species checklists with vouchers.
4. Other `R` packages provide tools to tackle some of the workflow steps described above. But in addition to the new features and resources related to the data editing and validation, the greatest strength of `plantR` is to provide a comprehensive and user-friendly workflow in one single environment, performing all tasks from data retrieval to export. Thus, `plantR` can help researchers better assess data quality and avoid data leakage in a wide variety of studies using species records.

## 1 INTRODUCTION

Biological collections (e.g. museums and herbaria) are essential for studying biodiversity (Graham et al., 2004). Taxonomists use these collections to describe new species, produce taxonomic revisions and species checklists, among other important uses (Funk, 2003; Bebber et al., 2010; Besnard et al., 2018). In macroecology, biogeography, and conservation, biological collections are often the main source of species records, which are used to study spatial patterns of biodiversity, species ecological niches, endemism levels, and conservation status (Graham et al., 2004; Dauby et al., 2017; Ulloa et al., 2017; Lima et al., 2020). Biological collections are increasingly making their electronic databases available in online databases, such as the Global Biodiversity Information Facility (GBIF). This growing availability of information has catalyzed many syntheses of our biodiversity knowledge (e.g. Antonelli et al. 2018), highlighting the importance of biological collections even more.

The increasing availability of biological collections databases has also exposed the wide variation of the documentation standards within and between collections (Willemse et al., 2008). Within collections, specimens collected by different people or in different periods may vary in their notation standards. The international documentation standards themselves are constantly evolving (www.tdwg.org/standards). Moreover, older records tend to have less associated information (e.g. missing geographical coordinates) and may contain names of localities that no longer exist (i.e. changing toponyms). Between collections, differences may emerge from different choices of documentation standards, on how to enter specimen information in the electronic databases, and on which fields should be entered first in the face of limited resources. The staff of biological collections often have little time to update the information that has been already entered in their databases or to correct data entry errors (e.g. typographical errors). These tasks become more challenging as the number of records in the collection increases.

Despite the global efforts to standardize the documentation of biodiversity information (e.g. Darwin Core standards), there is still much variation within fields associated with species records. This variation is likely to remain for years to come because biological collections are often underfunded, undervalued, and understaffed (de Gasper et al., 2020). Online databases, such as GBIF, gather, store, flag, and check some but not all the information provided by the data providers. This means that, although highly valuable, the available databases from biological collections are not always ready for use (Peterson et al., 2018). So, the final users of species records (e.g. taxonomists, ecologists, and conservationists) often have to decide between performing those procedures themselves or trusting the data available without knowing exactly the level of data quality. This is problematic because variation in data quality can impact the outcomes of studies in taxonomy, ecology, and conservation (Graham et al., 2004; Zizka et al., 2019; Rodrigues et al., 2020). Thus, we still need comprehensive and reproducible tools to manage species records from biological collections, particularly regarding notation standards, species identifications, duplicate records, and fine-scale validation of the geographical coordinates.

## 2 OVERVIEW

We present `plantR`, a new R package for managing species records from biological collections. As a general approach, `plantR` does not edit the original information; it stores the standardized information in new columns to assist collection curators in comparing original and edited information. Much of the new functionalities depend on gazetteers, maps, lists of taxonomists, and plant collections, which are provided with the package. As its name suggests, `plantR` was initially designed to manage plant records from herbaria, with some functionalities being currently exclusive to plants. However, if the input data has the required fields and data format, many `plantR` features should work for any group of organisms. `plantR` should interest taxonomists, biogeographers, ecologists, and conservationists, as well as curators of biological collections. The package is implemented in `R` (R Core Team, 2020) and details on its implementation and functionalities can be found at https://github.com/LimaRAF/plantR.

## 3 THE PLANTR WORKFLOW

`plantR` is accompanied by the proposal of a workflow to process the information associated with species records (Fig. 1). Here, we present the steps of this workflow and the main `plantR` features to apply it. They are presented in the order that the workflow should be applied. This order aims to maximize the edition and validation of the available information, although many `plantR` functionalities work independently from the previous steps of the workflow.

**FIGURE 1.**
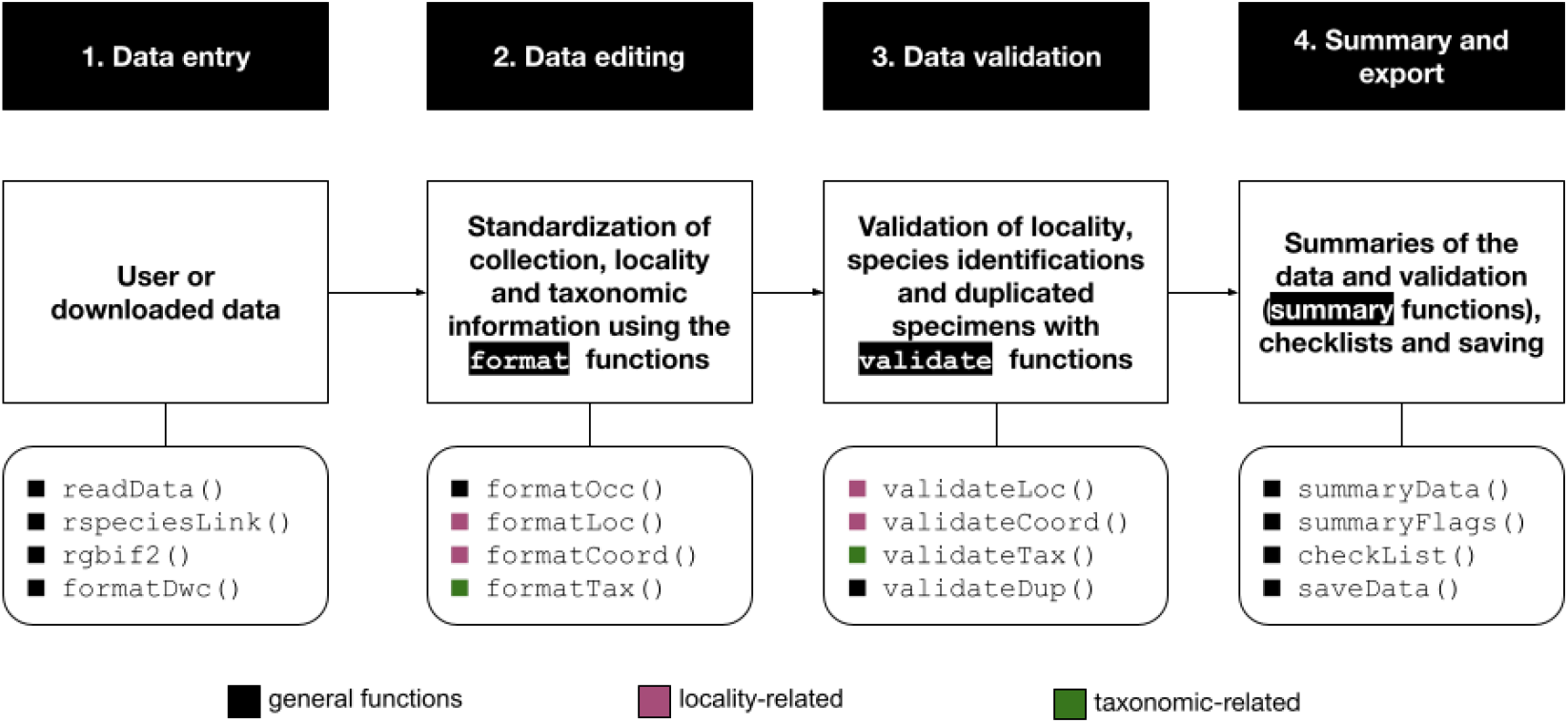
Chart illustrating the four main steps of the workflow proposed here to manage species records from biological collections for taxonomy, ecology, and biodiversity conservation. Black boxes represent each of the four steps, white boxes their description, and rounded boxes their main `plantR` functions.

### 3.1 Data entry

Users can download species records directly from R, which is currently done from the Centro de Refer-ência em Informação Ambiental (CRIA, www.cria.org.br) and GBIF (www.gbif.org), using functions `rspecieslink()` and `rgbif2()`, respectively. The function `rgbif2()` performs a search based on scientific names using the `rgbif` package, but with a standardized output to enter the `plantR` workflow. The function `rspeciesLink()` is more flexible allowing the user to search by scientific name or any other taxonomic level, collection, and locality. Since these two sources of species records return different fields, a function is provided to guarantee their correspondence with the DwC standards (function `formatDwc()`). Users can also load their own data, which can be converted to the Darwin Core (DwC) standards (https://dwc.tdwg.org) using the function `formatDwc()`. Alternatively, users can import data from zipped DwC-Archive files from a local directory or from a link for data download provided by GBIF (function `readData()`).

### 3.2 Data editing

Data standardization is particularly important when combining records from multiple collections, because they not always follow the same documentation standards. `plantR` provides tools to edit and standardize the notation of the information associated with the records, which are very important for validating locality information, assessing the confidence level of species identifications and searching duplicate records across collections (see 3.3 Data validation).

#### 3.2.1 People’s names and collection information

The first edits performed by `plantR` regards the name of collector and identifiers, collector’s number and collection year (function `formatOcc()`). By default, people’s names are returned in the Biodiversity Information Standards format (www.tdwg.org/standards/hispid3/), which is: last name + comma + initials separated by points (e.g. Gentry, A.H.). Name formatting takes into account generational suffixes (e.g. Junior), prepositions (e.g. da, dos, von), compound last names (e.g. Saint-Hilaire), some titles (e.g. Dr., Profa.) and multiple collector names. `plantR` also standardizes the collection codes using a database of over 5000 plant collection names and their respective Index Herbariorum or Index Xylariorum codes (function `getCode()`).

#### 3.2.2 Locality and spatial information

One of the innovations of `plantR` is the standardization of records’ locality information (i.e the DwC fields “country”, “stateProvince”, “municipality” and “locality”; function `formatLoc()`). For instance, names are transformed to English (e.g. Brasil or Brésil become Brazil) and their notation is standardized (e.g. BR or BRA become Brazil). In the case of missing locality information, `plantR` performs some text mining aiming to retrieve them from other fields. To make sure that the original or retrieved locality information does exist, the package cross-checks the locality information of records with a gazetteer (function `getLoc()`). This cross-checking is based on a standard name-string that hierarchically combines the locality information at the best resolution available, thus avoiding spurious matches of same locality names in different countries or states/provinces (function `strLoc()`). The default `plantR` gazetteer currently contains entries at country level for all countries and at the lowest administrative level available at GDAM (https://gadm.org) for all Latin American countries and dependent territories (e.g. U.S. Virgin Islands). For Brazil, the gazetteer also contains information at the locality level (e.g. farms, forest fragments, parks). Most importantly, users can provide their regional or personal gazetteers.

The gazetteer includes some of the most common spelling variants and historical changes to locality names (currently biased for Brazil), which allows collection curators to trace back the most up-to-date locality names to improve their databases (function `getAdmin()`). Additionally, `plantR` assigns a geographical coordinate from the gazetteer to all valid localities (function `getCoord()`), which can be used as working coordinates in the case of missing or problematic original coordinates. Besides the automated assignment of missing co-ordinates, the package formats the original geographical coordinates to obtain non-zero, non-missing coordinates in decimal degrees (function `prepCoord()`).

#### 3.2.3 Taxonomic information

`plantR` offers tools to format scientific name notation (function `fixSpecies()`), such as the isolation and removal of taxonomic rank (e.g. var., subsp.) and name modifiers (e.g. cf., aff.), which is important for records containing more raw taxonomic information (e.g. morpho-species, incomplete identifications). The package also standardizes the name of botanical families, using a list of valid family names and synonyms from the APG IV for angiosperms (Chase et al., 2016) and PPG I for lycophytes and ferns (Schuettpelz et al. 2016; function `prepFamily()`). If the family name is not found in the list, a search for a valid family name is performed based on the genus. Finally, the package can replace synonyms, orthographic variants and typographical errors in species names (function `prepSpecies()`, which is performed using functions from the packages `Taxonstand` (Cayuela et al., 2021) and `flora` (Carvalho, 2020). These packages perform exact and fuzzy name matching from The Plant List (www.theplantlist.org/) and the Brazilian Flora 2020 project (http://floradobrasil.jbrj.gov.br/), respectively.

### 3.3 Data validation

#### 3.3.1 Locality and spatial information

`plantR` compares the precision of the original locality information with the one obtained by the crosschecking with a gazetteer (function `validateLoc()`). This comparison allows to flag possible typographical errors or unknown place names, which users can drop from the analyses or double-check themselves depending on their goals. Obtaining valid locality information is essential for the validation of geographical coordinates because they are validated by comparing the locality information of the record and the locality obtained by overlapping the coordinates with administrative maps (function `checkCoord()`). The package offers procedures for detecting the inversion and/or swap of coordinates (function `checkInverted()`), coordinates falling in the sea or bays, near the shoreline (`checkShore()`), and in neighbouring countries (`checkBorders()`). If after these procedures the locality information from the record and maps matches, the coordinate is flagged as validated, with an indication of the resolution of the validation (i.e. country, state, municipality or locality levels). As before, the validation of geographical coordinates is done using maps at the country level for the world and at the lowest administrative level available at GDAM for Latin America, but users can provide their own maps. Finally, `plantR` also provides tools to detect records from cultivated individuals (function `getCult()`) and spatial outliers (function `checkOut()`), i.e. coordinates too far away from the core distributions for a given taxon (Liu et al., 2018).

#### 3.3.2 Species identifications

One highlight of `plantR` is the classification of records according to the confidence in their species identifications (function `validateTax()`). This validation is based on a global list of ca. 8500 plant taxonomists names compiled from different sources (Lima et al., 2020). By default, this classification assigns the highest confidence level to three different cases: (i) type specimens (e.g. iso-types, holotypes), (ii) records identified by a specialist of the family, and (iii) records collected by the specialist of the family but with the identifier field empty (case iii is optional). The confidence level of records without identifier information (including NA’s) is flagged as ‘unknown’, while records identified by non-family specialists it are flagged as ‘low’. Users can provide their own list of taxonomists, as long as this list has the same general format as the default list provided by `plantR`. More-over, `validateTax()` returns the most frequent names of identifiers that are not in the taxonomist list, allowing users to provide missing taxonomist names.

#### 3.3.3 Duplicate records

Another novelty of `plantR` regards duplicates, i.e. samples of the same specimen incorporated in two or more collections (function `validateDup()`). Sharing biological material across collections is a common and encouraged practice, and they can represent 25% or more of the records available for regional biotas (e.g. Lima et al., 2020). The search for duplicates in `plantR` is executed by combining fields related to the taxonomy, collection and locality of the records (e.g., family + collector name + collector number + municipality). Because of the great variation in the notation and completeness of collector’s and localities names, the package allows the simultaneous use of different combinations of these fields to search for duplicates (function `getDup()`). If two or more combinations are provided, the search of duplicates uses tools from network analysis to find both direct and indirect links between records. The retrieval of duplicates across collections performs well using relatively large data-sets (i.e. millions of records). However, finding all existing duplicates requires that the databases of all collections are available and that all search fields are complete and filled in without typos using the same notation standards (or notations that `plantR` can standardize). This is rarely the case, so the list of duplicates returned should be considered incomplete in many cases.

`plantR` provides not only tools to search for duplicates, but also to homogenize information within the groups of duplicates found, such as species, locality and/or spatial information (function `mergeDup()`). This homogenization allows retrieving the best information available within duplicates, which is particularly useful when collections vary in the number and completeness of the digitized fields. After this homogenization, users can choose to remove or not the duplicates from the data. See Lima et al. (2020) for more details on the search and merge of duplicates implemented here.

### 3.4 Data summary and export

As a final step of the workflow, `plantR` can help users to summarize their data (e.g. number of occurrences, collections and species; function `summaryData()`) and the flags of the validation process (i.e. localities, coordinates, identifications and duplicates; function `summaryFlags()`). The package also provides species checklists with user-defined numbers of voucher specimens and the export of records by groups (e.g. families, countries, collections).

## 4 IMPLEMENTATION

### 4.1 Example of usage

The `plantR` workflow can be implemented using few command lines and wrapper functions (see Table 1 for details). Here, we provide a simple example using only one species. A detailed tutorial of the package is provided at https://github.com/LimaRAF/plantR.

**TABLE 1.** List of the main functions per type of information and per step of the proposed workflow. We also present the wrappers of the main functions for each step (if present) and the other `R` packages necessary to execute them.

| Workflow step | Type of information | Main functions | Wrapper | Dependencies |
| --- | --- | --- | --- | --- |
| 1 - Data Entry | Species records | readData, rgbif2,<br>rspeciesLink,<br>formatDwc | - | rgbif,<br>data.table |
| 2 - Data Editing | Names, numbers, etc | prepName, colNumber,<br>getYear, getCode | formatOcc | stringr |
|  | Localities | fixLoc, strLoc,<br>prepLoc, getLoc | formatLoc | countrycode,<br>stringr |
|  | Coordinates | prepCoord, getCoord | formatCoord | - |
|  | Taxonomy | fixSpecies,<br>prepSpecies,<br>prepFamily | formatTax | flora,<br>Taxonstand,<br>data.table |
| 3 - Data Validation | Localities | validateLoc | - | - |
|  | Coordinates | checkCoord,<br>checkBorders,<br>checkShore,<br>checkInverted,<br>getCult, checkOut | validateCoord | sf, robustbase,<br>data.table |
|  | Species identification | validateTax | - | - |
|  | Duplicate records | prepDup, getDup,<br>mergeDup, rmDup | validateDup | data.table,<br>igraph |
| 4 - Summary and Export | Summaries | summaryData,<br>summaryFlags,<br>checklist | - | data.table,<br>stringr |
|  | Export | saveData | - | data.table |

~~~
*# Installing plant R*
remotes : : **install _** github (“ LimaRAF **/** plant R “)
**library** (“ plant R “)
*# Data download*
occs **_** splink **<**− rspecies Link (species = “ Euterpe edulis “)
occs **_** gbif **<**− rgbif2 (species = “ Euterpe edulis “)
occs **<**− formatDwc (splink **_ data** = occs **_** splink , gbif **_ data** = occs **_** gbif)
*# Data editing*
occs **<**− formatOcc (occs)
occs **<**− formatLoc (occs)
occs **<**− formatCoord (occs)
occs **<**− formatTax (occs)
*# Data validation*
occs **<**− validateLoc (occs)
occs **<**− validate Coord (occs)
occs **<**− validate Tax (occs)
occs **<**− validate Dup (occs)
*# Data summary*
summs **<**− summaryData (occs)
flags **<**− summaryFlags (occs)
checklist **<**− checkList (occs)
~~~

### 4.2 Dependencies on other packages

Some of `plantR`’s features depend on other `R` packages (Table 1). Function `rgbif2()` uses package `rgbif` (Chamberlain et al., 2021) for downloading GBIF data. The management of strings, countries names, and spatial data use packages `stringr` (Wickham, 2019), `countrycode` (Arel-Bundock et al., 2018), and `sf`, (Pebesma, 2018), respectively. As mentioned above, function `prepSpecies()` uses `Taxonstand` (Cayuela et al., 2021) and `flora` (Carvalho, 2020). The search of duplicates uses package `igraph` (Csardi and Nepusz, 2006) to perform indirect string search. Finally, many functions use `data.table` (Dowle and Srinivasan, 2020), which provides fast table manipulation, reading and saving.

## 5 DISCUSSION

### 5.1 Comparison with other R packages

Other `R` packages already provide spelling and synonym checks of species names (Chamberlain and Szöcs 2013; Cayuela et al. 2021; Carvalho 2020; Kindt 2020), so there was no need to ‘reinvent the wheel’ and their functionalities were (or will be) integrated in `plantR`. `CoordinateCleaner` (Zizka et al., 2019) provides a great toolbox to work with geographical coordinates and we suggest this package for the advanced editing of geographical coordinates. The differential of `plantR` lies in providing both locality and coordinate validation, the automatic retrieval of coordinates for missing or problematic coordinates, and the coordinate validation at the county level. However, because these validations depend on the package gazetteer, these innovations currently apply mainly to Latin America. `plantR` also provides an approach to find cultivated specimens (i.e. `getCult()`), which is based on the fields ‘locality’ or ‘occurrenceRemarks’ and thus different from the approach used by `CoordinateCleaner`.

We found only one package that validates species identifications, `naturaList` (Rodrigues et al., 2020). This package also uses the field ‘identifiedBy’, but it returns more confidence levels of species identification and requires a user-provided list of taxonomists. The differential of `plantR` relies on the provision of a large database of plant taxonomists, besides the possibility of the user providing an extra list of specialist names. In addition, `plantR` also relies on the field ‘typeStatus’ and it performs the validation at the family-level. We are not aware of other `R` packages that perform (i) the edition of people names, (ii) the validation of locality information and (iii) the search/merge of duplicates.

### 5.2 Limitations and future developments

The variation in the notation of names, numbers and dates associated with species records across biological collections is huge; `plantR` handles most but not all of them. We envisage having a dictionary of common collectors’ names, but today some double-checking is still necessary. As mentioned before, locality and county-level geographical validation are currently biased towards Latin America. Therefore, users must be aware that the package does not provide solutions to all problems related to species records information. Some improvements predicted to be implemented in the future include the download from other data repositories (e.g. JABOT, http://jabot.jbrj.gov.br), the expansion of the package gazetteer and county-level maps and the validation of species names against databases that have wider geographical and taxonomic coverage (e.g. Catalogue of Life). We also plan to include simple functions that prepare records to enter the workflow of other `R` packages (e.g. `modleR` or `ConR` - Sánchez-Tapia et al. 2020; Dauby et al. 2017), that facilitate the citation of collections (e.g. `occCite` - Owens et al. 2021) and that collect provenance (e.g. `rdt` - Lerner et al. 2018). Moreover, the gazetteer, list of taxonomists, maps, and collections are constantly being improved; we are open to receive and incorporate missing or regional information to make them more complete.

## 6 CONCLUDING REMARKS

The number of collection databases made available online has greatly increased in the last decades and will probably continue to increase in the years to come (Graham et al., 2004; Sweeney et al., 2018). Therefore, having tools to assess and improve the quality of the information associated with species record is a pressing issue in biodiversity research. `plantR` provides these tools, some of them being presented for the first time. Although there are packages that provide similar tools, the greatest strength of `plantR` is to provide a comprehensive toolbox and a user-friendly workflow to process species records from beginning to end within a single environment. Thus, we expect that `plantR` can improve the reproducibility of taxonomic, ecological and conservation studies. But more importantly, we hope that `plantR` can assist collection curators to flag possible issues that need attention, thus saving their time while conducting the important task of maintaining biological collections.

## ACKNOWLEDGEMENTS

This package was supported by the European Union’s Horizon 2020 research and innovation program under the Marie Skłodowska-Curie grant agreement No 795114. M.F.S., A.S.-T. and S.R.M. were supported by the Coordination for the Improvement of Higher Education Personnel - CAPES (process 88887.145924/2017-00), the PNPD/CAPES program and the PCI program of the ‘Instituto Nacional da Mata Atlântica’ (INMA), respectively. We thank Sidnei Souza from CRIA for his help with the web API. We also thank CNCFlora and the TreeCo database for providing localities used to construct the gazetteer, and Vinícius C. Souza (ESALQ/USP) who helped to curate the list of plant taxonomists.

## AUTHORS’ CONTRIBUTIONS

R.A.F.L. conceived the idea and R.A.F.L., A.S.-T., S.R.M. and M.F.S. designed methodology. R.A.F.L. constructed the list of taxonomists, collections, and families, while R.A.F.L., A.S.-T., S.R.M. constructed the gazetteer and maps. R.A.F.L., A.S.-T., S.R.M. and H.t.S. wrote the codes and package documentations. R.A.F.L. led the writing of the manuscript, with contributions from A.S.-T. All authors contributed critically to the manuscript and gave final approval for publication.

## DATA AVAILABILITY STATEMENT

The `R` package `plantR` is available at https://github.com/LimaRAF/plantR. The version of the package described in this paper (version 0.1.3) is archived at [link to be included before publication].

